# KmerStream: Streaming algorithms for *k*-mer abundance estimation

**DOI:** 10.1101/003962

**Authors:** Páll Melsted, Bjarni V. Halldórsson

## Abstract

**Motivation:** Several applications in bioinformatics, such as genome assemblers and error corrections methods, rely on counting and keeping track of *k*-mers (substrings of length *k*). Histograms of *k*-mer frequencies can give valuable insight into the underlying distribution and indicate the error rate and genome size sampled in the sequencing experiment.

**Results:** We present KmerStream, a streaming algorithm for computing statistics for high throughput sequencing data based on the frequency of *k*-mers. The algorithm runs in time linear in the size of the input and the space requirement are logarithmic in the size of the input. This very low space requirement allows us to deal with much larger datasets than previously presented algorithms. We derive a simple model that allows us to estimate the error rate of the sequencing experiment, as well as the genome size, using only the aggregate statistics reported by KmerStream and validate the accuracy on sequences from a PhiX control.

As an application we show how KmerStream can be used to compute the error rate of a DNA sequencing experiment. We run KmerStream on a set of 2656 whole genome sequenced individuals and compare the error rate to quality values reported by the sequencing equipment. We discover that while the quality values alone are largely reliable as a predictor of error rate, there is considerable variability in the error rates between sequencing runs, even when accounting for reported quality values.

**Availability:** The tool KmerStream is written in C++ and is released under a GPL license. It is freely available at https://github.com/pmelsted/KmerStream

*Contact:*

## 1 INTRODUCTION

*k*-mers are one of the most fundamental objects used when analyzing DNA sequencing data. Many assembly algorithms (Zerbino and Birney, 2008; Gnerre *et al.*, 2011; Li *et al.*, 2010) start by constructing a *de Bruijn* graph, a graph containing all *k*-mers for some fixed *k*. Some of the most commonly used algorithms for aligning reads to a reference genome start by finding short exact matches of a fixed length *k* (commonly referred to as a seed); an index of all *k*-mers is constructed and from this index an initial alignment of a part of the read is found.

Current methods for obtaining aggregate statistics of *k*-mer data are based on keeping track of all *k*-mers in a set of reads. Much work has been done on reducing memory requirements, based on exact or approximately correct methods of keeping track of a large set of *k*-mers. This work includes using succinct set representations (Conway and Bromage, 2011) or probabilistic encodings such as Bloom filters (Pell *et al.*, 2012; Chikhi and Rizk, 2012; Melsted and Pritchard, 2011). Although the impact on memory usage is considerable, compared to previous approaches, these methods require storing all *k*-mers, explicitly or implicitly, in memory. Thus the amount of resources will grow linearly with the input size. Many methods also rely on having access to all the reads for multiple passes over the data. Thus all of the above methods suffer from “the curse of deep sequencing” (Roberts *et al.*, 2011) in which more sequencing can overwhelm the program in terms of memory usage and the algorithms simply fail to make use of increased amounts of data.

We propose using streaming algorithms to solve this problem, a framework first proposed by Alon, Matias and Szegedy (Alon *et al.*, 1996). In this framework we assume that the algorithm has limited memory, compared to the size of the dataset, and data arrives one item at a time. The algorithm has limited time to process the item, usually constant time or logarithmic in the input, and cannot store all the data. Additionally, in the one pass setting, the algorithm can only observe each item once and can never look back into the stream. Several streaming algorithms have been discovered for a number of problems, such as counting the number of distinct items in the data stream using logarithmic memory (Bar-Yossef *et al.*, 2002), and finding heavy hitters (Cormode and Hadjieleftheriou, 2010), i.e. frequent items that occur more than a fixed percentage of the time. Most of these algorithms are approximate, and probabilistic, one can specify the degree of the approximation and the probability of correctness beforehand, which will in turn affect the memory requirements and running time. The bioinformatics community has not widely adopted streaming algorithms. We suspect this is in part because of the probabilistic requirement and the complexity of implementation. However, we argue that probabilistic algorithms are ideally suited for sequencing data because a) the sequencing experiment is inherently probabilistic, it is unlikely that the same dataset can be generated twice and b) if done properly the probability that the algorithms is correct can be taken as an input parameter, furthermore we can show that this probability is independent of the dataset.

Sequencing errors cause considerable problems for *k*-mer based algorithms for both assembly and read mapping; in assembly these may cause the assembly to become disconnected and increases both the memory usage and computational time. In read mapping it may lead to increased computational overhead and the true location of the read not being found. The removal or correction (Meacham *et al.*, 2011; Schröder *et al.*, 2009) of erroneous bases and erroneous fragments is therefore a common pre-processing step in the analysis of DNA sequences, in particular when doing de novo assembly or read mapping to a reference assembly. Several programs (Li *et al.*, 2010; Kelley *et al.*, 2010; Liu *et al.*, 2013) use *k*-mers to explicitly fix or remove sequencing errors in the dataset and it is recommended to use them prior to assembly (Salzberg *et al.*, 2012).

A number of software programs have been written for quality control of DNA sequencing data, including FastQC (Andrews, 2010). FastQC has a number of functionalities useful for quality control, including giving a distribution of the quality values assigned by the sequencer, quality distribution by position, N content and GC content, identifying overrepresented *k*-mers and sequence length distribution. However, the *k*-mers identified tend to be short, 6 basepairs by default, and although they are useful for identifying contaminants they are not unique enough in the underlying genome to be useful for assessing the sequencing error rate, independent of what is given by the DNA sequencers.

The main contribution of this paper is a streaming algorithm for estimating efficiently the number of *k*-mers that occur exactly once in a data set, taking care of identifying the *k*-mer with its reverse compliment. The time requirement for this algorithm is only a constant factor times the time taken to read the data and requires space that is only logarithmic in the size of the dataset. This method can be extended to give an estimate of the *k*-mer abundance histogram. Additionally our algorithm reports frequency moments of the *k*-mer count, which are aggregate statistics of the histogram.

For experimental validation we show the results of running KmerStream on reads from a single lane of PhiX control sequences. We show the distribution accuracy of the estimator compared to the accurate *k*-mer counts and that the estimators are almost always within the approximationi levels guaranteed. Additionally we a simple error model, both from the estimates KmerStream produces as well as the true *k*-mer counts. Both error estimates are then compared to the true *k*-mer error rate obtained from mapping *k*-mers to the reference genome of PhiX174. The results in section 4.1 show that our simple error model underestimates the true error rate, however only by a few percent on average.

As a simple application of our method we use it to estimate the the error rate in a sequencing dataset, conditioned on the per basepair quality value given by the instrument manufacturer. Unlike most other methods designed for estimating error of sequencing data, the method does not require the mapping of the data to a reference genome as a preprocessing step. The low computational overhead of our method makes it suitable to identify quality issues early on in the analysis pipeline. The method can therefore be used as a filtering step to determine the quality of a run, quickly determining errors before the read mapping or assembly stage. We ran the method on a set of 2656 whole genome sequenced individuals. Our results suggest that the error probabilities given by Illumina are largely accurate. Our results also show that once we have conditioned on an error probability given by the instrument manufacturer there is considerable variance in the sequencing error, suggesting that sequencing error needs to be reassessed on a per sample basis. We further construct a diploid genome containing all the called variants assigned to haplotypes and compute the number of *k*-mers in this diploid genome.

## 2 METHOD

We take as input a collection of short reads *S*. For each read *s ∈ S* we generate all substrings of length *k* in *s*, denoted as *k*-mers. When reads contain other characters than *ACGT*, such as *N*, we split the read on the non-*ACGT* characters and consider all *k*-mers in the split reads. For each *k*-mer we also consider the reverse complement of that *k*-mer, and in general make no distinction between the two when counting.

A common task is to generate the abundance histogram for the *k*-mers. We define *f_i_* to be the number of distinct *k*-mers that appear *i* times in the set of reads. Since modern sequencing technologies are rarely strand-specific, we process each *k*-mer and its reverse complement as if they were equal. The histogram is then simply a table of the *f_i_* values.

Generating the exact histogram requires storing a large number of *k*-mers in memory and can be done with *k*-mer-counting tools such as Jellyfish (Marçais and Kingsford, 2011) or BFCounter (Melsted and Pritchard, 2011). Counting all *k*-mers in high-throughput sequencing datasets requires tens, or even hundreds of gigabytes of memory, whereas the method we propose requires less than ten megabytes. Recently a method was proposed that generates an approximation of the histogram based on sampling *k*-mers (Chikhi and Medvedev, 2013), this method however does not have a guaranteed error rate associated with it.

We propose to not consider the exact histogram itself, but to compute statistics based on the histogram counts, which can be found using less memory and more speed than current methods by using streaming algorithms.

One of the key statistics we are interested in is *f*_1_, the number of *singleton k*-mers, i.e. *k*-mers that appear exactly once in the set of reads. Previous studies (Melsted and Pritchard, 2011) have shown that when a genome is being sequenced at relatively high coverage the majority of singleton *k*-mers do not come from the genome, and are generated from sequencing errors. Also, the number of singleton *k*-mers grows with increased coverage, whereas the number of *k*-mers from the genome will approach a fixed number as coverage increases.

Given the frequencies *f_i_* of the histogram, we define the *k*-th frequency moment, *F_k_*, as

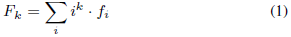

In addition to the number of singleton *k*-mers *f*_1_, we are interested in three moments; *F*_0_*, F*_1_ and *F*_2_. *F*_0_ is the number of distinct *k*-mers in the set of reads and *F*_1_ is simply the number of *k*-mers in the reads, counted with repetition. The second moment, *F*_2_, puts a higher weight on the number of *k*-mers that have high abundance values.

For each of the frequency moments streaming algorithms have been developed, that can estimate their value to within a factor of (1*±ε*) with high probability using only *O* (*ε*^−2^ · log(*N*)) memory, where *N* is the number of *k*-mers. It should be noted that estimating the frequency is solved in the general setting of counting arbitrary items in a stream, but for the remainder of the paper we will focus exclusively on the *k*-mer counting case.

Estimating the second moment, *F*_2_, was first solved in the seminal work of Alon, Matias and Szegedy (Alon *et al.*, 1996). This paper also formalized the framework and popularized the research field. The first rigorous estimator for the number of distinct elements, *F*_0_ is from (Bar-Yossef *et al.*, 2002), although earlier work from (Flajolet and Nigel Martin, 1985) applies as well. The first moment, *F*_1_, is easiest to construct as we can maintain a single counter that is incremented once for each *k*-mer.

### 2.1 Estimating *F*_0_ and *f*_1_

To compute the *f*_1_ estimator we use a hashing approach similar to the approach of Bar-Yossef et al. (Bar-Yossef *et al.*, 2002). The high level idea of the algorithm is to sample the stream at different rates, and afterwards select the sampling rate that gives the best result. For each *k*-mer *a* we compute the hash value *h*(*a*) and find the least significant 1 when the value is written in binary, e.g. if *h*(*a*) = 0110100_2_ then the least significant 1 is in the third position. For the zero value we define the position to be 64, or the number of bits used to represent the value. If the location of this bit is *w*, then 2*^w−^*^1^ divides *h*(*a*) and the binary representation of *h*(*a*) ends in 1, followed by *w* − 1 zeroes. Note that half the *k*-mers will have *w* = 1 a quarter will have *w* = 2, etc.

The data structure we use is a list of arrays, *T*, one for each potential value of *w*, we say that the array *T_w_* is at *level w*, and we say that all *k*-mers with value *w hash to this level*. Each array has a fixed size *R* that is only dependent on the error parameters *ε*, and each element in the array is a 2-bit number storing values from 0 to 3. When processing a *k*-mer *a* we look into the array *T_w_*. Conditioned on the value of *w* the *w* least significant bits are no longer random and so we discard the lowest *w* bits. We then use the result of the division, modulo the size of the array, an index into the array *T_w_* and increase the counter there. As we are only interested in *f*_1_, in the application presented here we limit the counter to two bits and if the counter is at 3, we do nothing. An extension of this work would be to allow the counter to count more bits, the algorithm could then be used to estimate *f_i_* for general *i*.

When the number of *k*-mers is much larger than *R* the array in *T*_1_ will almost certainly be full, i.e. all the values will be at 3. This indicates that sampling at a rate of one half is too small, and so we should look at lower sampling rates. Algorithm 1 estimates *f*_1_ by relating it to the probability that a counter in an array has value 0 or value 1 respectively. To get a good estimate for those probabilities we require that the number of distinct *k*-mers that hash to the array is roughly of the same size as the array itself. Thus after we have processed all the *k*-mers we can find the most appropriate array *T_w_* to use.

If we look at a fixed level *w*, the probability of a counter being 0 depends only on the number of distinct *k*-mers that hashed to this level, *N_w_*. This probability should be 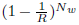, as the probability of each *k*-mer hashing to this counter is 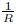. Note that the multiplicity of the *k*-mer does not factor in here, since each *k*-mer will always hash to the same counter every time it appears. If we can estimate this probability accurately, call it *p*_0_, we can solve for *N_w_* to get an approximation on the number of *k*-mers that hash to the level *w*. Note that solving

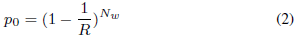

for *N_w_* yields

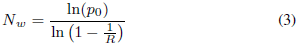

Since the sampling rate is 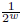 we can use this to approximate the number of distinct *k*-mers. Our estimator for *p*_0_ is 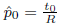 where *t*_0_ is the number of empty counters in the array *T_w_*. The estimator of the number of distinct *k*-mers is then

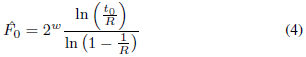

Although this would work for any value of *w*, i.e. all values of *w* have the same expectation, different values of *w* have different variance. If we use *w* = 1, then nearly all values in *T_w_* will be non-zero, giving a poor point estimate of *p*_0_. Similarly if *w* is too high, so that only a handful of elements map to *T_w_* the estimate for *p*_0_ will be poor. In general we want our estimates to be bounded away from 0 and 1, to this end we select the level *w*\* where the fraction of empty counters is as close to 50% as possible. By thinning the stream of hash values we can adaptively sample at different rates and decide on the best rate to use after we have seen all the data when we have to report the answer.

Up to this point we have followed the work of Bar-Yossef et al. (Bar-Yossef *et al.*, 2002), the estimator here is identical to the one derived in Theorem 2 of (Bar-Yossef *et al.*, 2002). We now show how to extend these results to obtain an accurate estimate of *f*_1_.

We define a singleton to be a *k*-mer that occurs once and let *x*_1_ be the number of singletons that hash to level *w*. We note that each counter with a value of 1 has to be the result of a singleton *k*-mer hashing to this counter, since for non-singleton *k*-mers the counter would have been increased at least twice. On the other hand some singleton *k*-mers will hash to the same counter as some other *k*-mer so we must find a way to relate *x*_1_ to some probability we can compute. The probability that a counter contains the value 1 is then

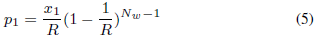

this is seen by choosing the singleton *k*-mer that should hash to the counter in *x*_1_ ways, it hashes to the counter with probability 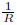 and all the other *Nw −*1 *k*-mers must not hash to the counter with probability 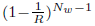. Note that from (2) we see that the part 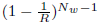 is equal to 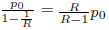. Thus we have 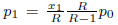, simplifying and solving for *x*_1_ yields

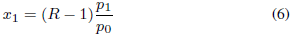

We estimate *p*_1_ in the same fashion as *p*_0_, namely let *t*_1_ be the number of counters in *T_w_* equal to 1 and set 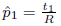.

We can expect the error in the estimates of the probabilities to be on the order of 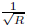, which will then translate to an error rate of similar magnitude for the *f*_1_ estimator.

#### Algorithm 1 Streaming algorithm for counting *k*-mers

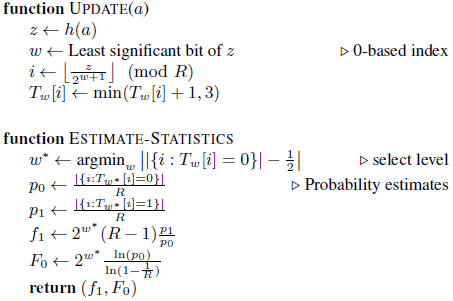

We formalize the selection of the optimal value of *w* in Theorem 1, the proof of this theorem can be found in the supplement for the paper.

#### THEOREM 1.

*If* 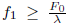 *then Algorithm 1 finds* 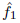 *such that* 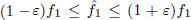 *with probability at least* (1 − *δ*). *The algorithm uses* 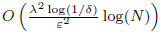 *memory and O*(1) *time per update, where N is the number of elements in the stream*.

Unlike for most statistical estimators, the theorem does not derive a distribution for the *f*_1_ statistic, but rather a bound on the probability that the point estimate deviates too far from the true value. In fact the estimator is slightly biased, since it uses the ratio of two unbiased estimators, *p*_0_ and *p*_1_ in Algorithm 1. However this bias is guaranteed to be smaller than the error rate *ε* used to select the parameter *R*.

We note that this method can be extended to obtain estimates of higher frequencies, *f_i_* for *i ≥* 2. As an example for *f*_2_, we note that if a counter has the value 2, this can only be obtained from two independent singletons mapping to the counter or one *k*-mer with frequency 2. The first happens with probability 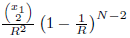 and the latter with probability 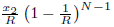, where *x*_1_ and *x*_2_ are the number of *k*-mers with frequency 1 and 2 that map to a level. Given an estimate for *N* and *x*_1_ we can solve to obtain an estimate for *x*_2_. This scheme could be generalized for higher values of *i* to obtain estimates of *f_i_*, although obtaining a guarantee on the error rate is left as an open problem.

### 2.2 Estimation of error rate

We can use our results to estimate error rates of a genomic sequencing experiment. For this we will focus on the *k*-mer error rate, i.e. given a *k*-mer from a read what is the probability that is did not originate from the DNA sequenced. This rate is higher than the basepair error rate, since we require all of the *k* basepairs to be intact. If the sequencing errors in a read were independently distributed, it would be easy to convert the *k*-mer error rate to a basepair error rate. However sequencing errors are not independent, generally the ends of the higher error rates and sequencing errors can come in batches. Thus we will focus on the *k*-mer error rate in this discussion and note that converting it to a basepair error rate assuming independence will lead to an underestimate of the true basepair error rate.

We use a simple generative model where total number of sampled *k*-mers with repetition, namely *F*_1_, is given. Each *k*-mer comes from a random position in the genome of size *G*, and with probability *ε_k_* it contains a sequencing error and is error-free with probability 1 − *ε_k_*. To account for repeated errors, we further assume that each erroneous *k*-mer is produced by sampling a random basepair and changing it to one of the three other nucleotides so that it does not match the reference. The number of *k*-mers sampled at each location follows a Poisson distribution with mean *λ* and so the coverage of an error-free *k*-mer is Poi(*λ*(1 *− ε_k_*). For a fixed erroneous *k*-mer the coverage follows a 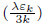, assuming that there is only one possible position in the genome that could produce this error. Our model has three parameters, *G, λ* and, *ε_k_*. To obtain estimates for these parameters we require three statistics, namely *f*_1_*, F*_1_ and, *F*_0_. Given the Poisson distributions of the coverages and the number of possible *k*-mers *G* for the error-free and *G* · 3 · *k* for the single error *k*-mers, we can derive the following equations

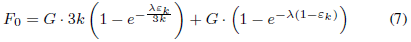

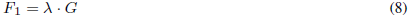

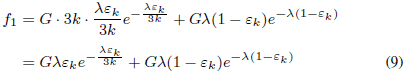

We can isolate *λ* from (8) and *G* from (7) to obtain

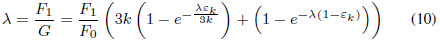

Similarly we derive by canceling the common factor *Gλ* from (9) using (8)

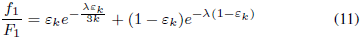

Using equations (10) and (11) we can solve for *λ* and *ε_k_* numerically to obtain an estimate of the coverage, *k*-mer error rate and finally the genome size.

We make a couple of observations about this model. First, the assumption that all error *k*-mers are solely due to single nucleotide differences causes us to slightly underestimate the true error rate. Second, a sequencing error in one *k*-mer may not be detected as an error because the error *k*-mer also occurs as a true *k*-mer in the genome. As we have chosen *k* = 31 then in random DNA the probability that a error *k*-mer also occurs in the genome is less than 10*^−^*^9^. Third, the same error *k*-mer can occur in two distinct reads by chance given high enough coverage, this is modeled in our *k*-mer distribution model via the Poisson variables. However the model does not take into account *k*-mers that have a high repeat count, due to repetitive regions or systematic errors in sequencing. According to our computations 96.6% of the genome of 31-mers in the genome occur exactly once or twice on a diploid genome. It has been observed that in practice sequencing reads are not randomly sampled from the genome and the true distribution is not a Poisson distribution. In particular the coverage of genomic regions is known to be dependent on the GC rate of the basepairs (Minoche *et al.*, 2011). Meacham et al. (Meacham *et al.*, 2011) have shown that the error rate is site specific, i.e. that sequencing error rate varies with the location being considered. Nonetheless we show in section 4.1 that this simplified model gives a good estimate of the *k*-mer error rate using only three statistics from the *k*-mer distribution, rather than the entire histogram.

## 3 COMPUTATIONAL EXPERIMENTS

### 3.1 Implementation

The software is implemented in C++ and can read FASTQ, compressed FASTQ and BAM files. The user can select the *k*-mer size used, the error rate *ε* and additionally it allows for filtering *k*-mers based on quality scores. For the quality filtering, all bases in a read that do not meet the cutoff are discarded, thus only *k*-mers where all the bases in the *k*-mer have good quality values are kept.

### 3.2 Setup

We ran our algorithm on a data set of 2656 whole genome sequenced individuals, using Illumina HiSeq sequencers. The individuals being sequenced had an average coverage of 15.9x and a minimum coverage of 6x. All of these data are sequenced under the same conditions at the same laboratory and have already undergone a number of quality control procedures (Styrkarsdottir *et al.*, 2013). We would expect these data to have comparable error characteristics.

All reads that were labeled as “qc-failure” or “optical or pcr duplicate” were discarded from our analysis. For the remaining reads we considered all *k*-mers that only contained basepairs that had a q-value above a fixed threshold. Hence, if a read contains a basepair with a q-score less than the given threshold only *k*-mers to the left and to the right of the basepair were considered. Our computations were run for q-score thresholds of 0, 13, 20 and 30, corresponding to basepair error rates of 1, 0.05, 0.01 and 0.001, respectively. Assuming independent error rates these basepair error rates should correspond to a lower bound of 100.0%, 79.6%, 26.8%, and 3.1% for the *k*-mer error rate, using the value *k* = 31. As an example, when using a q-score threshold of 30 then, according to the annotation given by the instrument manufacturer, all basepairs considered should have an error rate of less than 0.001 or 0.1%. As many of the basepairs will have even lower annotated error rates, then if the annotation of the manufacturer is correct, the average error rate should be even lower.

### 3.3 Counting *k*-mers in the human genome

We constructed the diploid genome of a single individual. As input we used Human reference genome build 36, genotypes called in that individual as determined using the GATK (DePristo *et al.*, 2011; McKenna *et al.*, 2010) genotype caller and the assignment of these genotypes to haplotypes using long range phasing (Kong *et al.*, 2008). From these we constructed two copies of each chromosome. We estimate a total of 2.552G 31-mers in this diploid genome, we observe that 4.9% of the 31-mers only occur once in the diploid genome, indicating that they overlap a polymorphic region, 91.7% of the 31-mers occur twice, indicating that the region is non-polymorphic, 0.1% occur three times, 1.9% of the 31-mers occur four times and 1.4% occur more than four times.

## 4 RESULTS

For running time and memory, we compare our software, KmerStream, to using KmerGenie on the H. Sapiens chr 14 (36M reads) and B. Impatiens (303 M reads), datasets from GAGE (Salzberg *et al.*, 2012). All tests were run on a Intel Xeon E-2650 16-core processors at 2.4 GHz with 128 Gb of memory. For the time comparison we only ran the histogram estimation step, the program specialk, and not the entire pipeline for KmerGenie.

It should be noted that the time to read the H. Sapiens dataset was 60 seconds and 1800 seconds for the compressed B. Impatiens dataset. Our experiments show that KmerStream is at least 2-3 times as fast as the previous approach and the memory usage is an order of magnitude better.

### 4.1 PhiX174 Validation

To validate the accuracy of the *k*-mer error rate model proposed in section 2.2 we used sequencing reads obtained from a PhiX control lane. The sequencing data from the control has 363M reads, while the genome size of PhiX174 is only 5386 nucleotides. Due to the small genome size, reads were sampled so that the per basepair coverage would be 30-fold. This was necessary to scale down the coverage to values the better correspond to normal sequencing coverage used for whole genome sequencing. This was repeated 1000 times and each computational experiment run independently. For each input we used *k* = 31 and classified all *k*-mers in reads as true or false, depending on whether they appeared in the reference sequence or not. Separate histograms were generated for both classes of *k*-mers. We then ran KmerStream for each input obtaining estimates for *f*_1_, *F*_1_ and *F*_0_. The top row of figure 1 shows the distribution of relative accuracy for *f*_1_ and *F*_0_ respectively. The program was run with a default error guarantee of 2% for *F*_0_, since for this input *f*_1_*/F*_0_ is about 0.5 this results in a 4% error guarantee of *f*_1_. From the distribution of figure 1 (top row), we see that the vast majority of the experiments have estimates within the error guaranteed.

**Fig. 1.**
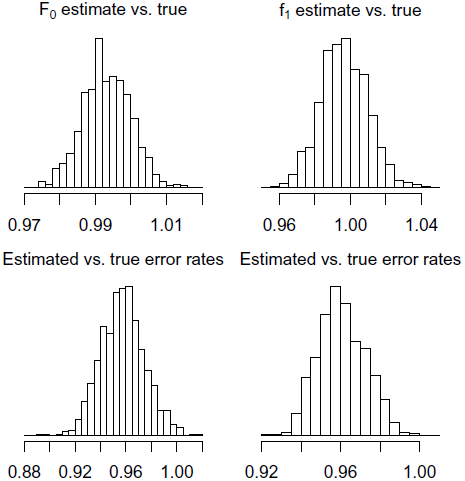
Relative accuracy of KmerStream. Top row: accuracy for *F*_0_ and *f*_1_ respectively. Bottom row: relative accuracy of *k*-mer error rate based on *k*-mer estimates (left) and true *k*-mer numbers (right).

The error rate model was fitted using both the true values for the statistics as well as the estimated values from KmerStream. The true *k*-mer error rate was obtained by matching *k*-mers to the PhiX174 reference genome.

Given the number of caveats listed when deriving our simplified model, as well as obtaining estimates from only three key statistics, we are pleasantly surprised to see how well the model fits the actual results. Based on the true values for the *k*-mer statistics we see from figure 1 (bottom row, right) that our model underestimates the *k*-mer error rate by 4% on average. When using the estimated values from KmerStream, figure 1 (bottom row, left) shows that we underestimate the error rate by 4% on average, same as using the true *k*-mer statistics, but we observe an increase in the variance of the accuracy of our estimate. This increase in variance is primarily because of the underlying variance in our estimate of the *k*-mer statistics. Of course the KmerStream method is much more efficient than obtaining the accurate counts, both in terms of runtime and memory.

### 4.2 Error rates

To estimate the error rates we ran a set of experiments on a set of 2656 whole genome sequenced individuals. Figure 2 shows a histogram of the average per *k*-mer error rate for different q-value thresholds and in Table 2 we give critical values of the distribution of the read error rate. We observe that without any filtering on q-values, on average 5.9% of the *k*-mers are estimated to contain errors. This number varies considerably between samples and 5% of the samples have an estimated error rate of 11.5% or higher, while 5% of the samples have an estimated error rate of 3.7% or lower.

**Fig. 2.**
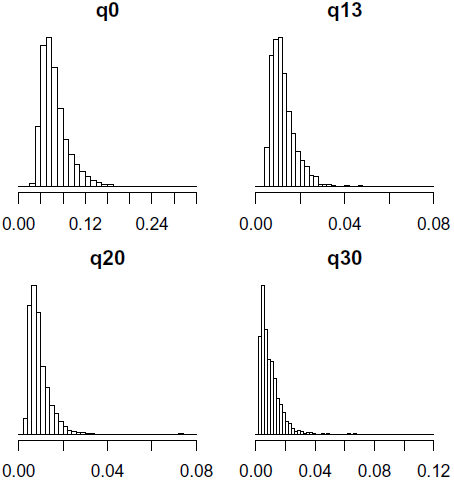
*k*-mer error rates for different quality filters for 2656 individuals.

**Table 1.** Software comparison for running time and memory usage.

| Software | Parameter (k) | Dataset | Time | Memory |
| --- | --- | --- | --- | --- |
| KmerStream | 21 | H. Sapiens, chr14 | **194s** | **11M** |
| KmerGenie | 21 | H. Sapiens, chr14 | 580s | 127M |
| KmerStream | 31 | H. Sapiens, chr14 | **220s** | **11M** |
| KmerGenie | 31 | H. Sapiens, chr14 | 703s | 126M |
| KmerStream | 31 | B. Impatiens | **3120s** | **11M** |
| KmerGenie | 31 | B. Impatiens | 7620s | 114M |

**Table 2.** Percentiles of *k*-mer error rates at different q-value thresholds for 2656 individuals.

| Threshold | Percentile |  |  |
| --- | --- | --- | --- |
|  | 5 | 50 | 95 |
| q0 | 3.7% | 5.9% | 11.4% |
| q13 | 0.6% | 1.1% | 2.2% |
| q20 | 0.4% | 0.8% | 1.8% |
| q30 | 0.3% | 0.8% | 2.2% |

We observe that the error rate decreases with a higher q-value threshold. We observe a marked uniform decline in the fraction of error *k*-mers when the threshold is increased from 0 to 13, an average reduction of 80%.. A smaller, but clear, decrease is observed when the q-value threshold is increased from 13 to 20, 25% on average. However when comparing increasing the threshold from 20 to 30 we observed an average change of −0.9%, but a median of 9%. Additionally this decrease in the number of error *k*-mers is not uniform and in 44% of all individuals the estimated error rate increases when the q-value threshold is increased from 20 to 30, i.e. using q-value thresholds larger than 20 does not necessarily give one lower error rate *k*-mers. Furthermore the number of *k*-mers, i.e. the coverage of the data set was reduced by 25% on average by increasing the q-value threshold from 20 to 30.

In Figure 2 we consider only the *k*-mers that are removed when the error threshold is increased, giving a histogram of the fraction of those that are error *k*-mers. We observe that when increasing the q-value threshold from 0 to 13, on average an estimated 35% of the *k*-mers removed are error *k*-mers. This suggests that unless the algorithm being used for analysis is highly robust to errors then using a q-value threshold lower than 13 will in many cases be problematic. On average of 85% of all *k*-mers in our dataset have q-value at all bases greater than 13, indicating that not considering *k*-mers with a q-value threshold less than 13 has limited impact on the number of *k*-mers being considered while it has a large impact on the fraction of *k*-mers that are error free. When we increase the q-value threshold from 13 to 20 an average of 5% of the *k*-mers removed are error *k*-mers and when increasing the threshold from 20 to 30 an average of 1.5% of the *k*-mers removed are error *k*-mers.

## 5 DISCUSSION

The amount of data being gathered with modern sequencing methods continues to grow at a faster rate than our ability to analyze and store the data. An alternative view to the current state of the art is to consider technologies that sequence DNA “on the fly”. In this case the sequencing machine does not store all the results, but rather transmits the sequence reads as they are generated. Technologies that fit this framework have been proposed (Branton *et al.*, 2008; Clarke *et al.*, 2009) and are currently in development, such as technologies from Oxford Nanopore, but technical details are limited at this point in time. Regardless of the exact technologies used, this new sequencing paradigm opens up new opportunities for online or streaming analysis of the data, where we bypass the storage requirements, and simply plug the sequencing directly into the analysis. The upshot of developing streaming algorithms for analysis of genetics datasets is that they are not only efficient in terms of computational resources, but also future-proof in terms of new sequencing paradigms.

One benefit of the algorithms we have developed is as follows; that *F*_0_ − *f*_1_ is a crude estimate of the number of *k*-mers that have been sequenced at least twice, when this number goes above a certain fraction of the genome size we can decide to stop sequencing. Another benefit is that when the error rate goes above some threshold we can decide to stop the experiment immediately, not wasting our time on failed experiments. The method presented here can be particularly useful when used for a species that has not been previously sequenced, allowing us to get an estimate the coverage of this genome while sequencing prior to assembly.

When we condition on the error rate given by Illumina we see considerable variability in the error rate between individuals. Hence, it is not advisable to use the error rates in a model without considering differences between individuals.

Our results show that although the base pair quality values given by the instrument manufacturer are largely correct, there appears to be a considerable sample dependent difference in the the error rate conditioned on the base pair quality rate reported by the manufacturer. Our recommendation based on the results of sequencing 2656 individuals is to estimate both the number of *k*-mers *F*_0_ as well as the coverage and *k*-mer error rate for multiple q-value thresholds and decide on a case by case basis.

## REFERENCES

Alon, N., Matias, Y., and Szegedy, M. (1996). The space complexity of approximating the frequency moments. In Proceedings of the twenty-eighth annual ACM symposium on Theory of computing, pages 20–29. ACM.

Andrews, S. (2010). FastQC A Quality Control tool for High Throughput Sequence Data. http://www.bioinformatics.babraham.ac.uk/projects/fastqc/.

Bar-Yossef, Z., Jayram, T., Kumar, R., Sivakumar, D., and Trevisan, L. (2002). Counting distinct elements in a data stream. In Randomization and Approximation Techniques in Computer Science, pages 1–10. Springer.

Branton, D., Deamer, D. W., Marziali, A., Bayley, H., Benner, S. A., Butler, T., Di Ventra, M., Garaj, S., Hibbs, A., Huang, X., et al. (2008). The potential and challenges of nanopore sequencing. Nature biotechnology, 26(10), 1146–1153.

Chikhi, R. and Medvedev, P. (2013). Informed and automated k-mer size selection for genome assembly. Bioinformatics.

Chikhi, R. and Rizk, G. (2012). Space-efficient and exact de Bruijn graph representation based on a Bloom filter. In Algorithms in Bioinformatics, pages 236–248. Springer.

Clarke, J., Wu, H.-C., Jayasinghe, L., Patel, A., Reid, S., and Bayley, H. (2009). Continuous base identification for single-molecule nanopore DNA sequencing. Nature nanotechnology, 4(4), 265–270.

Conway, T. C. and Bromage, A. J. (2011). Succinct data structures for assembling large genomes. Bioinformatics, 27(4), 479–486.

Cormode, G. and Hadjieleftheriou, M. (2010). Methods for finding frequent items in data streams. The VLDB Journal, 19(1), 3–20.

DePristo, M. et al. (2011). A framework for variation discovery and genotyping using next-generation DNA sequencing data. Nature Genetics, 43(5), 491–8.

Flajolet, P. and Nigel Martin, G. (1985). Probabilistic counting algorithms for data base applications. Journal of computer and system sciences, 31(2), 182–209.

Gnerre, S., MacCallum, I., Przybylski, D., Ribeiro, F. J., Burton, J. N., Walker, B. J., Sharpe, T., Hall, G., Shea, T. P., Sykes, S., et al. (2011). High-quality draft assemblies of mammalian genomes from massively parallel sequence data. Proceedings of the National Academy of Sciences, 108(4), 1513–1518.

Kelley, D., Schatz, M., and Salzberg, S. (2010). Quake: quality-aware detection and correction of sequencing errors. Genome Biology, 11(11), R116.

Kong, A., Masson, G., Frigge, M. L., Gylfason, A., Zusmanovich, P., Thorleifsson, G., Olason, P. I., Ingason, A., Steinberg, S., Rafnar, T., et al. (2008). Detection of sharing by descent, long-range phasing and haplotype imputation. Nature genetics, 40(9), 1068–1075.

Li, R., Zhu, H., Ruan, J., Qian, W., Fang, X., Shi, Z., Li, Y., Li, S., Shan, G., Kristiansen, K., Li, S., Yang, H., Wang, J., and Wang, J. (2010). De novo assembly of human genomes with massively parallel short read sequencing. Genome Research, 20(2), 265–272.

Liu, Y., Schröder, J., and Schmidt, B. (2013). Musket: a multistage k-mer spectrum-based error corrector for Illumina sequence data. Bioinformatics, 29(3), 308–315.

Marçais, G. and Kingsford, C. (2011). A fast, lock-free approach for efficient parallel counting of occurrences of k-mers. Bioinformatics, 27(6), 764–770.

McKenna, A., Hanna, M., Banks, E., Sivachenko, A., Cibulskis, K., Kernytsky, A., Garimella, K., Altshuler, D., Gabriel, S., Daly, M., et al. (2010). The Genome Analysis Toolkit: a mapreduce framework for analyzing next-generation DNA sequencing data. Genome research, 20(9), 1297–1303.

Meacham, F., Boffelli, D., Dhahbi, J., Martin, D., Singer, M., and Pachter, L. (2011). Identification and correction of systematic error in high-throughput sequence data. BMC bioinformatics, 12(451), 1–11.

Melsted, P. and Pritchard, J. (2011). Efficient counting of k-mers in DNA sequences using a Bloom filter. BMC Bioinformatics, 12(1), 333.

Minoche, A. E., Dohm, J. C., Himmelbauer, H., et al. (2011). Evaluation of genomic high-throughput sequencing data generated on illumina hiseq and genome analyzer systems. Genome Biol, 12(11), R112.

Pell, J., Hintze, A., Canino-Koning, R., Howe, A., Tiedje, J. M., and Brown, C. T. (2012). Scaling metagenome sequence assembly with probabilistic de Bruijn graphs. Proceedings of the National Academy of Sciences, 109(33), 13272–13277.

Roberts, A., Pachter, L., et al. (2011). RNA-Seq and find: entering the RNA deep field. Genome medicine, 3(11), 74.

Salzberg, S. L., Phillippy, A. M., Zimin, A., Puiu, D., Magoc, T., Koren, S., Treangen, T. J., Schatz, M. C., Delcher, A. L., Roberts, M., et al. (2012). GAGE: A critical evaluation of genome assemblies and assembly algorithms. Genome Research, 22(3), 557–567.

Schröder, J., Schröder, H., Puglisi, S. J., Sinha, R., and Schmidt, B. (2009). SHREC: a short-read error correction method. Bioinformatics, 25(17), 2157–2163.

Styrkarsdottir, U., Thorleifsson, G., Sulem, P., Gudbjartsson, D. F., Sigurdsson, A., Jonasdottir, A., Jonasdottir, A., Oddsson, A., Helgason, A., Magnusson, O. T., et al. (2013). Nonsense mutation in the LGR4 gene is associated with several human diseases and other traits. Nature.

Zerbino, D. R. and Birney, E. (2008). Velvet: algorithms for de novo short read assembly using de Bruijn graphs. Genome research, 18(5), 821–829.

